# Methamphetamine Conditioned Place Preference in Adolescent Mice: Interaction Between Sex and Strain

**DOI:** 10.1101/2025.11.17.688853

**Authors:** Lewis Nunez Severino, Andre B. Toussaint, Rachael M. Langa, Grace McKenna, Isabella Bodziony, Nesha S. Burghardt

## Abstract

**Background:** Methamphetamine (METH) use is higher in adolescent women than men. While rodent studies support a sex difference in the reinforcing effects of METH, few have investigated sex differences in the underlying neural circuits, none of which tested rodents during adolescence.

**Aims:** Investigate whether there are sex differences in the rewarding effects of METH in two strains of adolescent mice that are commonly used to generate transgenic lines. Identify changes in associated neural circuits.

**Methods:** We tested METH-induced conditioned place preference (CPP) in male and female 129Sv/Ev and C57Bl/6 mice during middle adolescence using 1mg/kg of METH. Behaviorally-induced upregulation of c-Fos protein expression was quantified in the nucleus accumbens (NAc) and area CA1 of the hippocampus following the post-conditioning test.

**Results:** In C57Bl/6 mice, METH induced CPP in females, but not males. Conversely, METH-induced CPP in 129Sv/Ev males, but not females. In both strains, groups that exhibited CPP had more c-Fos+ cells in the NAc and CA1 when compared to saline-treated control groups. The number of c-Fos+ cells in these two brain regions correlated in groups exhibiting CPP, indicating increased NAc-CA1 communication during retrieval of the conditioned memory. Finally, we found evidence of behavioral sensitization in 129Sv/Ev males only.

**Conclusions:** Our study reveals that the rewarding effects of the METH in adolescent mice are both sex- and strain-dependent, indicating that METH response may result from an interaction between sex-specific and genetic mechanisms. Our findings will be informative when selecting an appropriate background strain in future studies using genetically modified mice.

## Introduction

Addiction to methamphetamine (METH) continues to be a public health crisis, currently affecting approximately 1.6 million people in the United States (Jones et al., 2020). METH use in adolescents is on the rise, resulting in a 20% increase in emergency room visits by individuals 12-17 years of age (Administration, 2022; Administration, 2023). This is particularly concerning given that drug taking during adolescence increases the risk of developing a lifelong substance use disorder (Chambers et al., 2003; Grant and Dawson, 1998; Jordan and Andersen, 2017), which may be attributed, in part, to the lasting effects of drugs on the developing adolescent brain. In addition, the biological response to psychostimulants during adolescence tends to be more rewarding and less aversive, potentially increasing drug taking behavior (Schramm-Sapyta et al., 2009). However, the majority of preclinical studies investigating the neural circuits mediating the rewarding effects of METH have used adult rodents (Liu et al., 2021; Wang et al., 2023; Su et al., 2022; Yao et al., 2024) and effects of METH on the adolescent brain are understudied.

Interestingly, there are sex differences in the use and response to METH in humans. Adolescent females are more likely to use METH than adolescent males (Rawson et al., 2005). Women also begin use earlier and have a shorter transition to regular use than men (Dluzen and Liu, 2008; Brecht et al., 2004; Hser et al., 2005). While both sexes are polysubstance users, women are more likely to report METH as their main drug of choice (Cretzmeyer et al., 2003). Men and women also initiate use for different reasons. Women often begin using METH for weight loss purposes, to increase productivity, and to treat symptoms of depression, while men frequently report initiating use because their parents use drugs and for recreational purposes (Cretzmeyer et al., 2003; Brecht et al., 2004). Sex differences in METH intake have also been found in rodents, with female rats exhibiting greater self-administration and faster escalation of METH intake than males (Kucerova et al., 2009; Roth and Carroll, 2004; Reichel et al., 2012). While these findings are suggestive of sex differences in the reinforcing effects of METH, few studies have investigated sex differences in the underlying neural circuits, none of which administered METH to rodents during adolescence.

Sex disparities in addiction are often attributed to sex hormones, but genetic mechanisms may also play a role in these effects (Krueger et al., 2023). Genetic factors are known to contribute importantly to individual differences in susceptibility to addiction (Bierut, 2011; Hiroi and Agatsuma, 2005). Similarly, genetic variation in rodents affects behavioral response to drugs of abuse, as evidenced by strain differences in the effects of METH, alcohol, cocaine, amphetamine, and phencyclidine on locomotor behavior (Alexander et al., 1996; Phillips et al., 2008; Schlussman et al., 1998; Good and Radcliffe, 2011). While there are numerous studies characterizing behavioral differences among mouse strains, they frequently involve only one sex and interactions between hormonal and genetic mechanisms are rarely considered. Identification of strain differences in response to METH in both sexes would be informative when selecting an appropriate background strain for studies targeting underlying mechanisms in transgenic mice.

Here, we used conditioned place preference (CPP) to test the rewarding effects of METH in male and female adolescent mice of two inbred strains, both of which are commonly used for generating genetically modified mice (Barnabei et al., 2010; Seong et al., 2004). To evaluate the neural basis of METH-induced CPP, we quantified behaviorally-induced upregulation of c-Fos protein expression in the nucleus accumbens (NAc) and area CA1 of the hippocampus. We found that the effects of METH on behavioral sensitization, CPP, and neural activity in C57Bl/6 and 129Sv/Ev mice are both strain- and sex-dependent. This work indicates that sex and genetics interact to affect response to drugs of abuse and that changes in communication between the NAc and hippocampus may contribute importantly to the rewarding effects of METH in the adolescent brain.

## Materials and methods

### Animals

Male and female C57Bl/6 and 129Sv/Ev mice were purchased from Taconic Biosciences (Germantown, NY) at postnatal day (PND) 21 and group housed (4/cage) with mice of the same sex. They were maintained on a 12-hour light/dark cycle (05:00 -17:00) with free access to food and water. Experiments began on PND 40 (middle adolescence) (Laviola et al., 2003) and were performed during the light cycle, with each sex tested separately. Procedures were approved by the Institutional Animal Care and Use Committee of Hunter College and were conducted in accordance with the National Institutes of Health Guidelines on the Care and Use of Laboratory Animals.

### Drug

Methamphetamine hydrochloride (Sigma-Aldrich, St. Louis, MO) was dissolved in 0.9% sterile saline and injected intraperitoneally (i.p.) at a dose of 1 mg/kg. It was made fresh daily prior to administration. Animals were weighed immediately before each injection to ensure the accuracy of drug dosage given.

### Apparatus

Conditioned place preference was conducted in a two-compartment apparatus separated by a removable Plexiglas partition (Columbus Instruments, Columbus, OH). One compartment (20.3 x 30.5 x 30.5 cm) contained white walls and floors and was cleaned with orange Clorox wipes (light compartment). The other compartment was the same size but had black walls and a red floor with strips of white tape that provided texture. It was cleaned with 100% ethanol (dark compartment). A small clear holding chamber (17.1 x 7.9 x 12.1 cm) provided access to both compartments when the adjoining door was opened.

### Conditioned Place Preference (CPP) Testing

The CPP procedure included preconditioning (day 1), conditioning (days 2-9), and a postconditioning test session (day 10) (**Fig. 1a**). Prior to the start of CPP, mice were habituated to handling (2 minutes) and adapted to the experimental room for two consecutive days (PND 38-39). During preconditioning (PND 40), mice were briefly placed in a holding chamber where they had access to both sides of the CPP box. Upon leaving the holding chamber, mice were allowed to freely explore the 2-chamber apparatus for 30 minutes (day 1). This was followed by a conditioning phase (PND 41-48), during which mice in the drug group received METH (1 mg/kg, i.p.) and were confined to the light compartment for 30 minutes every other day for 8 days (days 2, 4, 6, 8). On alternate days, the drug group received a saline injection before being confined to the dark compartment for 30 minutes (days 3, 5, 7, 9). Mice in the saline group received saline before being confined to either the light (days 2, 4, 6, 8) or dark (days 3, 5, 7, 9) compartment for 30 minutes. We paired drug with the light compartment, because we found that both strains exhibited an innate preference for the dark compartment during preconditioning (**Supplementary Fig. 1**). Pairing drug with the light compartment therefore improved our ability to detect increases in time spent on the drug-paired side after conditioning. Finally, there was a drug-free postconditioning test (PND 49), during which mice were again placed in the holding chamber before being given access to both compartments for 30 minutes. An overhead camera recorded behavior during preconditioning, the first and last days of drug administration, and the postconditioning test session. Videos were later analyzed by investigators blind to treatment condition using ANY-maze software (Stoelting, Wood Dale), which tracked each mouse and calculated distance traveled (m) and time in each compartment. A CPP score was calculated as the difference in time spent within the light compartment (i.e., drug-paired compartment) during pre- and postconditioning (postconditioning time – preconditioning time), with positive numbers indicating stronger preference for the light compartment after conditioning.

**Fig. 1.**
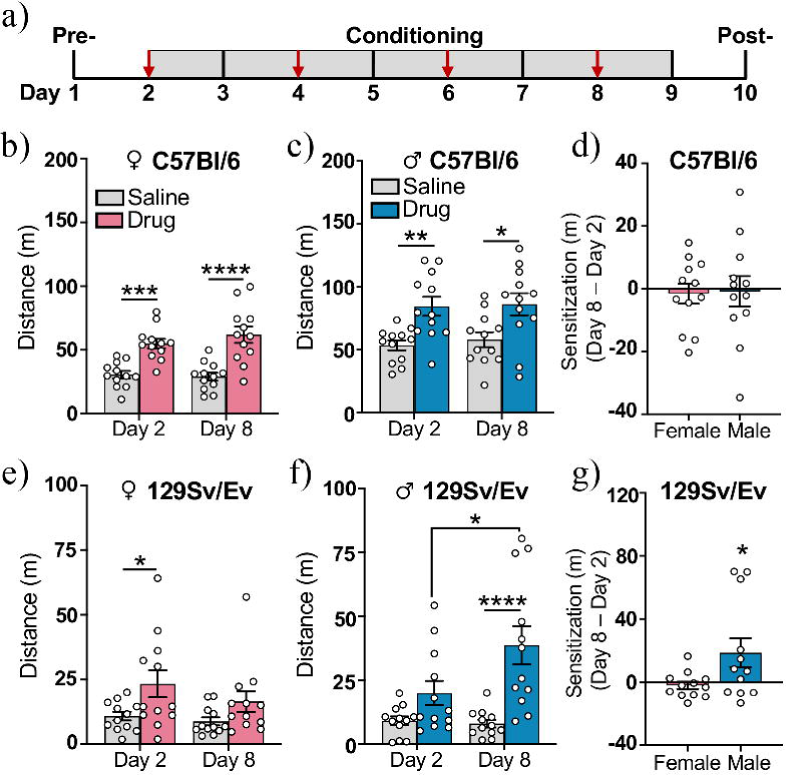
Methamphetamine-induced stimulation of locomotor activity. **(A)** Schematic of behavioral procedures. Red arrows indicate days that methamphetamine was administered. **(B-C)** Distance traveled following an injection (i.p.) of either methamphetamine (drug) or saline on the first day (experimental day 2) and the last day (experimental day 8) of treatment for C57Bl/6 **(B)** females and **(C)** males**. (D)** The difference in distance traveled during the first and last day of drug administration (meters traveled on day 8 – day 2) in drug-treated C57Bl/6 females and males. **(E-F)** Distance traveled following an injection (i.p.) of either methamphetamine (drug) or saline in 129Sv/Ev **(E)** females and **(F)** males on the first (day 2) and last day (day 8) of drug-treatment**. (G)** The difference in distance traveled during the first and last day of drug administration (meters traveled on day 8 – day 2) in drug-treated 129Sv/Ev females and males. C57Bl/6 females (n=12/group), C57Bl/6 males (n=12/group), 129Sv/Ev females (n=12/group), 129Sv/Ev males (n=12/group). Data represent mean ± SEM. *p<0.05, **p<0.01, ***p<0.001, ****p<0.0001.

### C-Fos Immunohistochemistry

Ninety minutes after the postconditioning test, a subset of mice were deeply anesthetized with a mixture of ketamine (100 mg/kg, i.p.) and xylazine (7 mg/kg, i.p.) and transcardially perfused with cold PBS followed by cold 4% paraformaldehyde in PBS. Brains were removed, postfixed in 4% paraformaldehyde overnight at 4-8°C, and then cryoprotected in 30% sucrose with 0.1% NaN_3_ for at least a week. Brains were sectioned into 35 mm thick coronal slices on a cryostat (CM3050S, Leica, Germany) and stored in PBS containing 0.1% NaN_3_ until staining. Every 6^th^ section containing the nucleus accumbens (NAc) and hippocampus was processed for c-Fos immunoreactivity. Free-floating sections were washed in PBS-0.1% triton (PBST) (3x, 10min) before being treated with 3% H_2_O_2_/50% methanol in PBST (15min). Tissue was then blocked with 10% normal donkey serum in PBST at room temperature (2 hours) before it was incubated in primary antibody (rabbit anti-c-Fos, 1:5,000; Santa Cruz Biotechnology, sc-52) in 10% normal donkey serum in PBST overnight at 4-8°C. The next day, tissue was washed in PBS (3x, 10min) and then incubated in a biotinylated secondary antibody (donkey anti-rabbit, 1:500; Jackson ImmunoResearch) in 10% normal donkey serum for 2 hours at room temperature. Tissue was then washed in PBS (3x), incubated in an avidin-biotin-peroxidase complex (ABC Elite Kit, Vector Laboratories) for 1 hour, and then visualized with 3,3’-diaminobenzidine (DAB Substrate Kit, Vector Laboratories).

### Cell Quantification

Every sixth section containing the NAc and the entire dorsoventral extent of the hippocampus was imaged at 10x using an upright Olympus BX53 light microscope. Immunoreactive cells were manually quantified by an experimenter blind to group who used Image-J software (Version 2.14, NIH) to create regions of interest (ROI) around the NAc core, NAc shell, and the CA1 subregion of the hippocampus. Four sections containing the NAc (+1.42 to +1.10 mm from bregma), five sections containing dorsal CA1 (-1.58 to -2.06 mm from bregma), and five sections containing ventral CA1 (-2.92 to -3.40 mm from bregma) were included in the analysis. Cells within each subregion were counted bilaterally and averaged across sections.

### Statistical Analysis

Data were analyzed with Student’s *t*-tests for independent samples, Pearson’s Correlation Coefficient, and two-way ANOVA, followed by Bonferroni’s post-hoc test when significant main effects or interactions were detected. All statistical analyses were performed using GraphPad Prism software (GraphPad, San Diego, CA) and significance levels were set at p<0.05.

## Results

### Innate preference for the dark compartment is stronger in 129Sv/Ev than C57Bl/6 mice

We first investigated whether mice had an innate preference for one side of the CPP box by quantifying time spent in each side (light vs. dark) during preconditioning. In the C57Bl/6 strain, a two-way repeated measures ANOVA revealed no sex x compartment interaction (F_(1,46)_ = 1.88, p = 0.18), but a significant effect of compartment (F_(1,46)_ = 101.7, p<0.0001) (**Supplementary Fig. 1a**). Similarly, the results of the analysis for 129Sv/Ev mice revealed no sex x compartment interaction (F_(1,46)_ = 5.83e-007, p = 0.99), but a significant effect of compartment (F_(1,46)_ = 3534, p<0.0001) (**Supplementary Fig. 1b**), demonstrating that mice of both strains and sexes preferred the dark compartment. We next evaluated whether there is a strain difference in the magnitude of this effect by calculating the difference in time each mouse spent in each compartment (dark time -light time). Consistent with previous findings indicating that 129 substrains have higher levels of anxiety (Rodgers et al., 2002), the two-way ANOVA revealed a significant main effect of strain (F_(1,92)_ = 52.64, p<0.0001), with 129Sv/Ev mice of both sexes exhibiting a stronger preference for the dark compartment than C57Bl/6 mice of both sexes (**Supplementary Fig. 1c**). These findings informed our decision to pair METH with the light compartment, as it allowed us to detect increases in time spent on the drug-paired side after conditioning.

### Methamphetamine induces behavioral sensitization in 129Sv/Ev males only

Repeated exposure to METH has been shown to induce behavioral sensitization in adult male C57Bl/6 mice, as measured by enhanced stimulation of locomotion over time (Liu et al., 2019; Li et al., 2021). Here, we investigated whether administering METH to adolescent mice every other day for 4 days leads to behavioral sensitization in a sex or strain-dependent manner. Distance traveled (meters) during the first and last day of METH administration (experimental days 2 and 8) was compared in saline and drug-treated groups (**Fig. 1a**). In C57Bl/6 females, a two-way repeated-measures ANOVA revealed no drug group x day interaction (F_(1,22)_ = 1.02, p = 0.32), but a significant effect of drug group (F_(1,22)_ = 45.25, p<0.0001), with Bonferroni’s post-hoc comparisons revealing that METH increased locomotion similarly on both days (day 2, p<0.001; day 8, p<0.0001) (**Fig.1b**). In C57Bl/6 males, there was also no drug group x day interaction (F_(1,22)_ = 0.15, p = 0.71), but a significant effect of drug group (F_(1,22)_ = 11.04, p<0.01), with METH increasing locomotion similarly on both days (day 2, p<0.01; day 8, p<0.05) (**Fig. 1c**). To directly compare behavioral sensitization in C57Bl/6 mice of both sexes, we calculated the difference in distance traveled from the first to the last day of drug administration (day 8 – day 2) in drug-treated mice only. We found that the mean difference was slightly below zero in both sexes (females: x □= -1.61 ± 3.21; males: x□= -0.82 ± 4.86) with no difference between the sexes (t_(22)_ = 0.14, p = 0.89) (**Fig. 1d**). Together, these results indicate that METH in C57Bl/6 mice stimulated movement on both days of administration, but there was no evidence of behavioral sensitization in either sex.

Interestingly, METH affected locomotion differently in 129Sv/Ev mice. In 129Sv/Ev females, a two-way repeated-measures ANOVA revealed no drug group x day interaction (F_(1,22)_ = 0.81, p = 0.38), but a significant effect of drug group (F_(1,22)_ = 5.87, p < 0.05) that was driven by drug-induced increases in locomotion on the first day of administration (day 2, p < 0.05) (**Fig.1e**). In contrast, the two-way repeated-measures ANOVA in 129Sv/Ev males revealed a significant drug group x day interaction (F_(1,22)_ = 4.41, p < 0.05), a significant effect of drug group (F_(1,22)_ = 21.85, p < 0.001), and an effect of day that approached significance (F_(1,22)_ = 3.49, p = 0.075). While there was no effect of METH on the first day (p = 0.20), METH significantly increased locomotor activity on the last day of administration (p<0.0001) (**Fig.1f**). Our direct comparison of drug-treated 129Sv/Ev males and females on measures of behavioral sensitization (distance traveled on day 8 – day 2) revealed that over time, METH elicited a significantly bigger increase in locomotion in males (x□= +18.67 ± 9.10) than females (x□= -2.02 ± 2.35) (t_(22)_ = 2.20, p < 0.05) (**Fig.1g**). Collectively, these results indicate that adolescent C57Bl/6 mice of both sexes are initially more sensitive to the stimulating effects of METH on locomotion than adolescent 129Sv/Ev mice, but that repeated exposure to METH leads to behavioral sensitization in 129Sv/Ev males only.

### Methamphetamine induces conditioning place preference in C57Bl/6 females and 129Sv/Ev males

To evaluate the rewarding effects of METH, we calculated a CPP score for each animal (time in light chamber postconditioning – preconditioning) and compared drug-and saline-treated groups. In C57Bl/6 mice, drug-treated females had significantly higher CPP scores than saline-treated females (t_(22)_ = 2.51, p < 0.05) (**Fig. 2a)**, while C57Bl/6 males of both groups had similar CPP scores (t_(22)_ = 0.03, p = 0.97) (**Fig 2b**). When we normalized CPP scores of drug-treated animals to same-sex control groups and directly compared the sexes, we found that C57Bl/6 females had significantly higher CPP scores than C57Bl/6 males (t_(22)_ = 5.70, p<0.0001) (**Fig. 2c**).

**Fig. 2.**
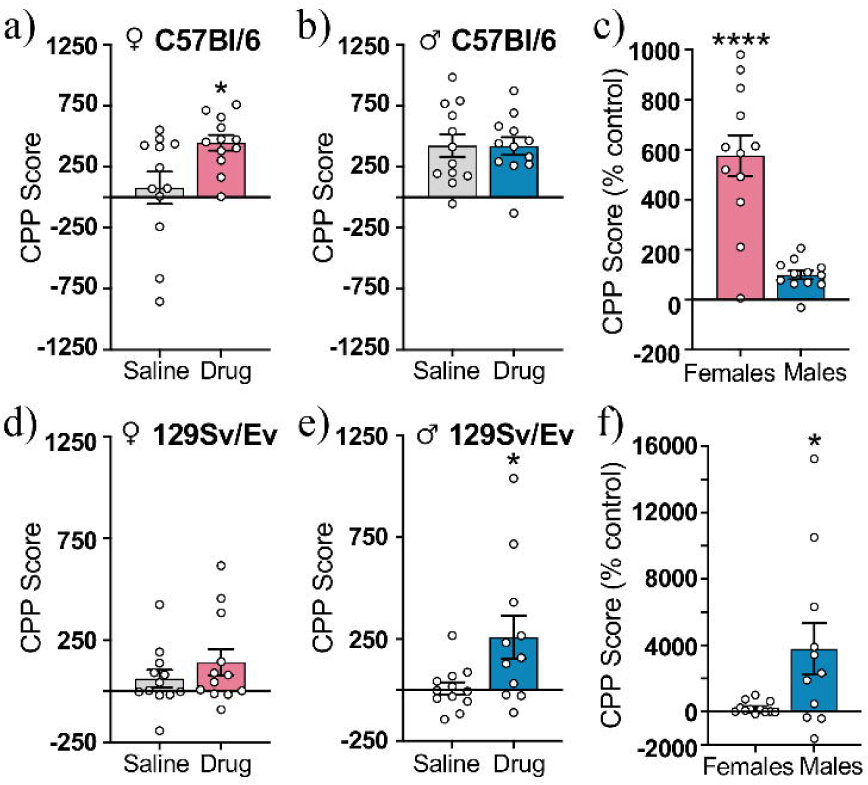
Methamphetamine-induced CPP in adolescent C57Bl/6 and 129Sv/Ev mice of both sexes. The CPP scores of C57Bl/6 **(A)** females and **(B)** males. **(C)** CPP scores of drug-treated C57Bl/6 females and males calculated as a percentage of same-sex saline-treated control groups. CPP scores of 129Sv/Ev **(D)** females and **(E)** males. **(F)** CPP scores of drug-treated 129Sv/Ev females and males calculated as a percentage of same-sex saline-treated control groups. C57Bl/6 females (n=12/group), C57Bl/6 males (n=12/group), 129Sv/Ev females (n=12/group), 129Sv/Ev males (n=11-12/group). Data represent mean ± SEM. ****p<0.0001, *p < 0.05.

We found a different pattern of results in 129Sv/Ev mice. Saline- and drug-treated 129Sv/Ev females had similar CPP scores (t_(22)_ = 1.05, p = 0.31) (**Fig. 2d**), but drug-treated 129Sv/Ev males had significantly higher CPP scores than saline-treated males (t_(21)_ = 2.38, p < 0.05) (**Fig. 2e**). When we normalized the CPP scores of drug-treated animals to same-sex control groups, 129Sv/Ev males had significantly higher CPP scores than 129Sv/Ev females (t_(21)_ = 2.40, p < 0.05). (**Fig. 2f**). Together, these results demonstrate that METH selectively induced CPP in adolescent C57Bl/6 females and adolescent 129Sv/Ev males, indicating that there are sex differences in the rewarding effects of METH that vary by mouse strain.

### Methamphetamine-induced CPP is associated with increased neural activity in the NAc

To investigate the neural correlates of METH-induced CPP, we collected brains following the postconditioning test and quantified behaviorally-induced upregulation of c-Fos in the NAc core and shell (**Fig. 3a**). In the C57Bl/6 strain, drug-treated females had more total c-Fos+ cells in the NAc than saline-treated females (t_(8)_ = 2.38, p < 0.05) (**Fig. 3b, left**). When the NAc core and shell were analyzed separately, the two-way repeated measures ANOVA revealed a significant effect of drug group (F_(1,8)_ = 5.68, p < 0.05), with post-hoc tests indicating that METH-treated females had more c-Fos+ cells specifically in the NAc shell than saline-treated controls (p < 0.05) (**Fig. 3b, right**). In contrast, previous exposure to METH in C57Bl/6 males did not affect the total number of c-Fos+ cells in the NAc (t_(8)_ = 0.97, p = 0.36) (**Fig. 3c, left**) or the number of labeled cells in either the NAc core or shell (drug group, F_(1,8)_ = 0.94, p = 0.36; core, p = 0.14; shell, p >0.99) (**Fig. 3c, right).** Interestingly, CPP testing upregulated c-Fos+ cells more in the NAc shell than the NAc core in both sexes, but this was less pronounced in C57Bl/6 males (effect of subregion, F_(1,8)_ = 10.49, p < 0.05) than C57Bl/6 females (effect of subregion, F_(1,8)_ = 259.5, p<0.0001).

**Fig. 3.**
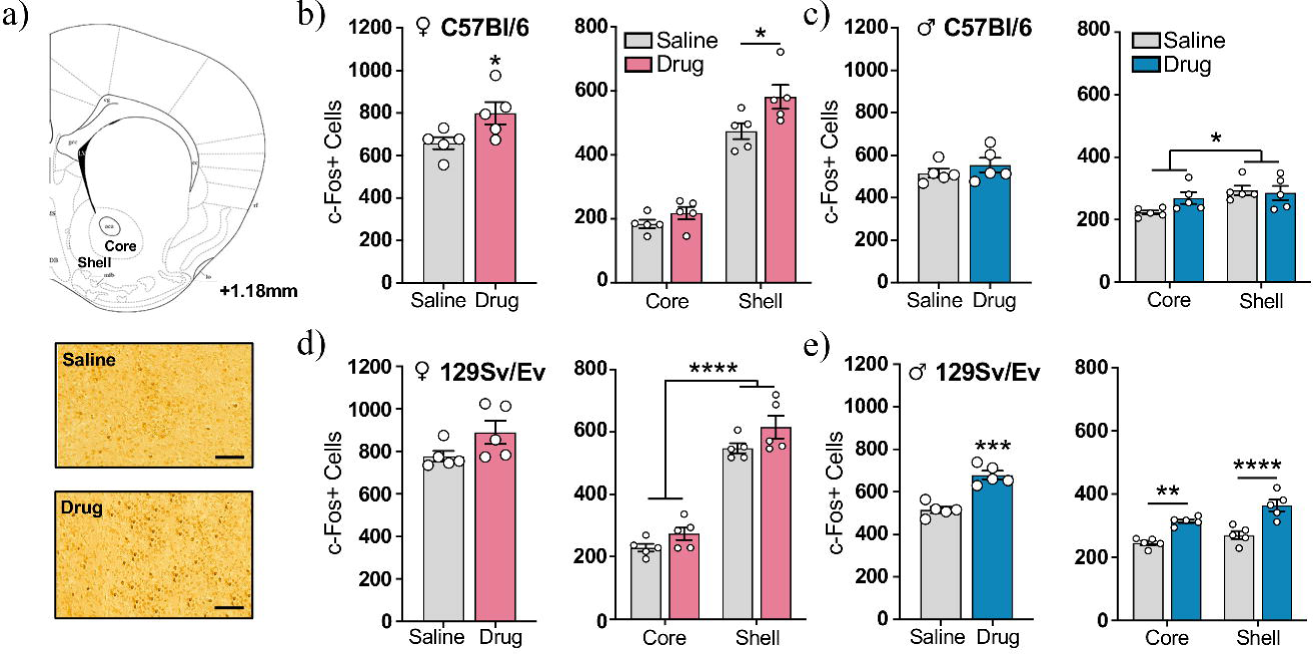
Behaviorally-induced upregulation of c-Fos in the NAc during the postconditioning test. **(A)** (Top) Schematic of a coronal section containing the NAc core and shell. (Bottom) Representative photomicrograph of c-Fos immunoreactivity in the NAc shell of a saline-treated and drug-treated C57Bl/6 female mouse. Scale bars, 100μm. **(B-C)** (Left) Total number of c-Fos+ cells in the NAc and (Right) number of c-Fos+ cells in each NAc subregion in C57Bl/6 **(B)** females and **(C)** males of each treatment group. **(D-E)** (Left) Total number of c-Fos+ cells in the NAc and (Right) number of c-Fos+ cells in each NAc subregion in 129Sv/Ev **(D)** females and **(E)** males of each treatment group. C57Bl/6 females (n=5/group), C57Bl/6 males (n=5/group), 129Sv/Ev females (n=5/group), 129Sv/Ev males (n=5/group). Data represent mean ± SEM. *p < 0.05, ** p < 0.01, ***p <0.001, ****p < 0.0001.

In the 129Sv/Ev strain, the two-way repeated measures ANOVA on females revealed no group differences in the total number of c-Fos+ cells in the NAc (t_(8)_ = 1.89, p = 0.10) or the number of c-Fos+ cells in either NAc subregion (drug group, F_(1,8)_ = 3.56, p = 0.10; core, p = 0.37; shell, p = 0.11) (**Fig. 3d**). In contrast, previous exposure to METH in 129Sv/Ev males significantly increased the total number of c-Fos+ cells in the NAc (t_(8)_ = 6.52, p < 0.001) (**Fig. 3e, left**), which was attributed to increases in both NAc subregions (drug group, F_(1,8)_ = 42.46, p < 0.001; core, p < 0.01; shell, p<0.0001) (**Fig. 3e, right**). Similar to the C57Bl/6 strain, CPP testing in 129Sv/Ev mice upregulated c-Fos+ cells more in the NAc shell than the NAc core, but this effect was again less pronounced in males (effect of subregion, F_(1,8)_ = 10.59, p < 0.05) than females (effect of subregion, F_(1,8)_ = 599, p<0.0001). These results demonstrate that the groups that exhibited METH-induced CPP had increased neural activity in the NAc during the postconditioning test.

### Methamphetamine-induced CPP is associated with increased neural activity in CA1

The hippocampus is a brain region that communicates directly with the NAc (LeGates et al., 2018; Barnstedt et al., 2024) and is critical for memory retrieval after conditioning (Kim and Fanselow, 1992). To gain further insight into the circuit mediating METH-induced CPP in adolescent mice, we expanded our c-Fos analysis to include area CA1 of the hippocampus, a brain region that has been implicated in CPP in previous studies (Assar et al., 2016; Rezayof et al., 2007). Our analysis included the dorsal and ventral poles of CA1, which differ both anatomically and functionally (Tao et al., 2021; Fanselow and Dong, 2010; Igarashi et al., 2014) (**Fig. 4a**). In C57Bl/6 mice, we found that drug-treated females had more total c-Fos+ cells in the CA1 than saline-treated controls (t_(8)_ = 3.33, p < 0.05) (**Fig. 4b, left**). METH increased c-Fos in both poles (two-way repeated measures ANOVA, drug group, F_(1,8)_ = 11.08, p < 0.05), but post-hoc tests revealed that this only approached significance in dorsal CA1 (p = 0.08) and reached significance in ventral CA1 (p < 0.05) (**Fig. 4b, right).** In contrast, we detected no group differences in c-Fos+ cells in the CA1 of C57Bl/6 males (total CA1, t_(8)_ = 1.72, p = 0.12; drug group, F_(1,8)_ = 2.94, p = 0.12; dorsal pole, p = 0.63; ventral pole, p = 0.17) (**Fig. 4c**) or 129Sv/Ev females (total CA1, t_(8)_ = 0.33, p = 0.75; drug group, F_(1,8)_ = 0.11, p = 0.75; dorsal pole, p>0.99; ventral pole, p>0.99) (**Fig. 4d**). However, drug-treated 129Sv/Ev males had more total c-Fos+ cells in the CA1 than saline-treated controls (t_(8)_ = 4.28, p < 0.01), effects that were attributed to increases in both the dorsal and ventral poles of the hippocampus (two-way repeated measures ANOVA, drug group, F_(1,8)_ = 18.32, p < 0.01; dorsal pole, p < 0.05; ventral pole, p < 0.01) (**Fig. 4e**). Together, these findings indicate that mice that exhibited METH-induced CPP had increased neural activity in the CA1 during the postconditioning test, results that mirror what we found in the NAc.

**Fig. 4.**
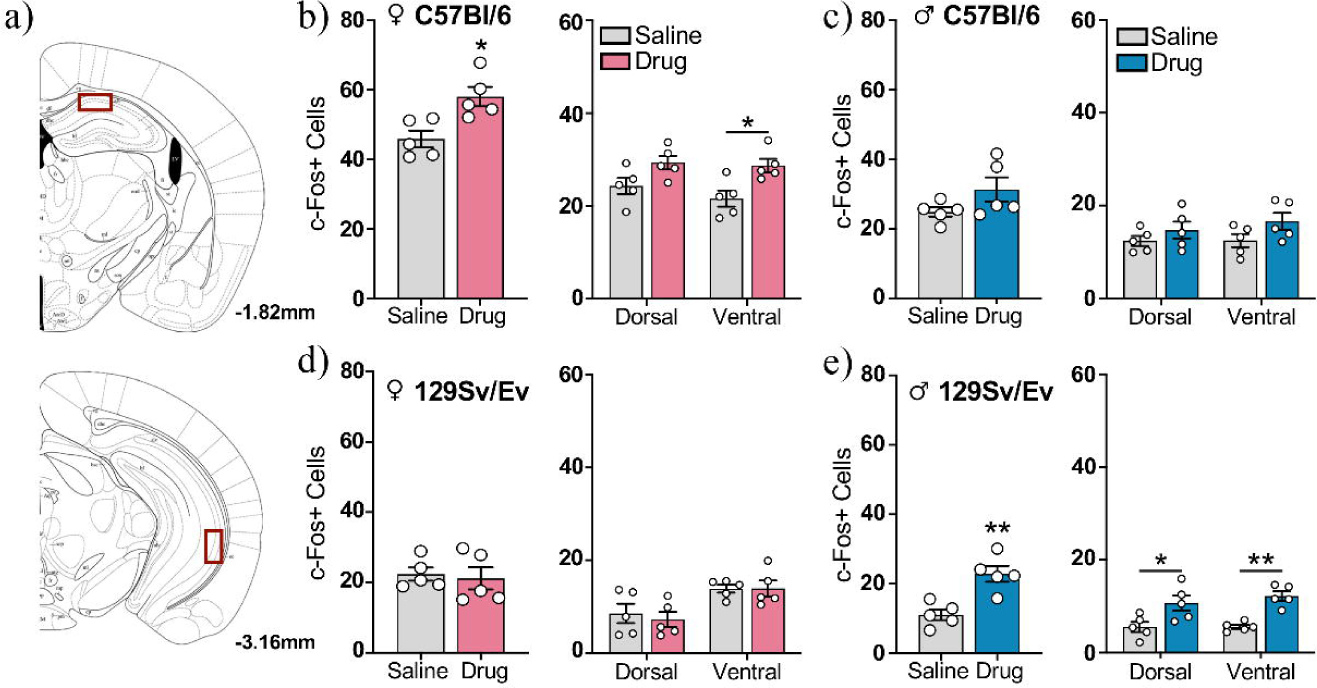
Behaviorally-induced upregulation of c-Fos in the CA1 subregion of the hippocampus during the postconditioning test. **(A)** Schematic of a coronal section containing the (Top) dorsal and (Bottom) ventral CA1 subregion of the hippocampus. The red box indicates the region used for cell quantification. **(B-C)** (Left) Total number of c-Fos+ cells in the CA1 and (Right) number of c-Fos+ cells in the dorsal and ventral poles of the CA1 in C57Bl/6 **(B)** females and **(C)** males of each treatment group. **(D-E) (**Left) Total number of c-Fos+ cells in the CA1 and (Right) number of c-Fos+ cells in the dorsal and ventral poles of the CA1 in 129Sv/Ev **(D)** females and **(E)** males of each treatment group. C57Bl/6 females (n=5/group), C57Bl/6 males (n=5/group), 129Sv/Ev females (n=5/group), 129Sv/Ev males (n=5/group). Data represents mean ± SEM. *p < 0,05, **p < 0.01.

### Neural activity in the NAc and CA1 is correlated during methamphetamine-induced CPP

To evaluate whether the NAc and hippocampus are part of the same circuit mediating METH-induced CPP, we examined whether their neural activity is correlated during the postconditioning test. In C57Bl/6 mice, we found that the total number of c-Fos+ cells in the NAc was positively correlated with the total number of c-Fos+ cells in the CA1 in females (r = 0.59, p < 0.01) (**Fig. 5a**) but not males (r = 0.20, p = 0.19) (**Fig. 5b**). In 129Sv/Ev mice, we found the same positive correlation between c-Fos+ cells in the NAc and CA1 in males (r = 0.65, p < 0.001) (**Fig. 5d**), but not females (r = 0.04, p = 0.59) (**Fig. 5c**). These findings demonstrate that neural activity was only correlated in mice that exhibited CPP and indicate that METH affects communication between the NAc and CA1 in a sex- and strain-specific way.

**Fig. 5.**
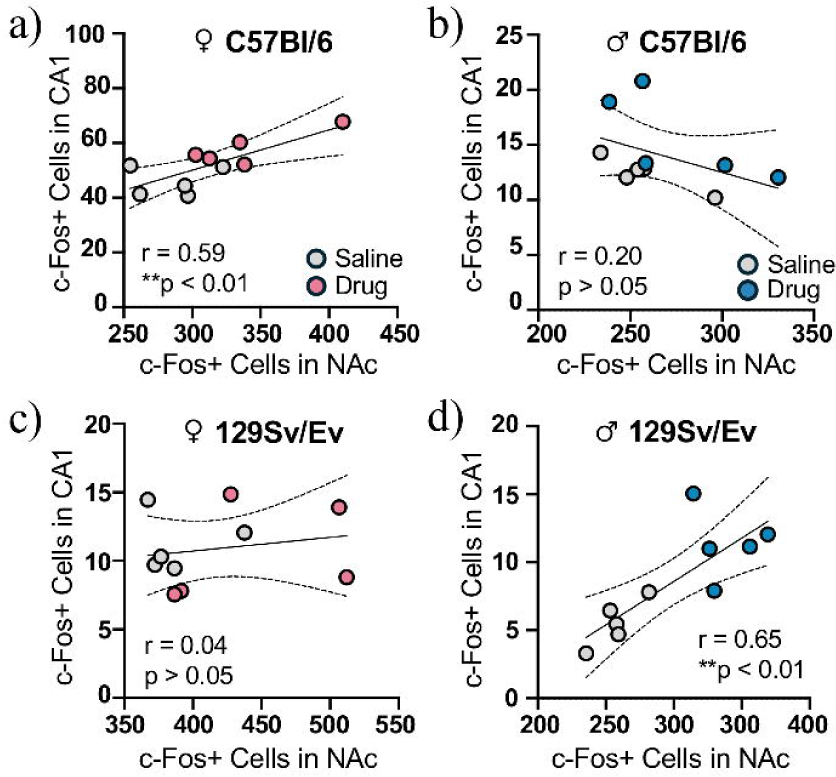
Correlated c-Fos expression in the NAc and CA1 subregion of the hippocampus. **(A-D)** Correlation between the number of c-Fos+ cells in the NAc and the number of c-Fos+ cells in the CA1 in drug- and saline-treated **(A)** C57Bl/6 females, **(B)** C57Bl/6 males, **(C)** 129Sv/Ev females and **(D)** 129Sv/Ev males. C57Bl/6 females (n=5/group), C57Bl/6 males (n=5/group), 129Sv/Ev females (n=5/group), 129Sv/Ev males (n=5/group), **p<001.

## Discussion

We tested the effects of METH on CPP in adolescent male and female mice of two different strains. In the C57Bl/6 strain, we found that a low dose of METH (1mg/kg) induced CPP in female, but not male mice. Conversely, in the 129Sv/Ev strain, we found that the same dose of METH induced CPP in male, but not female mice. Our analysis of c-Fos+ cells following the postconditioning test revealed that METH-induced CPP was associated with increased neural activity in the NAc and area CA1 of the hippocampus and increased communication between these two brain regions. These findings demonstrate that the rewarding effects of METH in adolescent mice are both sex- and strain-dependent and indicate that sex-specific biological mechanisms and genetics can interact to affect response to drugs of abuse. The sex and strain differences we report in wild-type mice will be informative for future CPP studies that target underlying mechanisms in transgenic mice and require selection of an appropriate background strain.

Numerous previous studies have tested the effects of METH on CPP in C57Bl/6 mice, as this is the most commonly used inbred mouse strain in biomedical research (Deacon et al., 2007). In contrast to our findings in adolescent males of this strain, METH has been shown to induce CPP in adult C57Bl/6 males following administration of the same dose (1mg/kg) (Chen et al., 2003; Liang et al., 2023; Shang et al., 2022), a higher dose (2mg/kg) (Lominac et al., 2016; Li et al., 2023; Szumlinski et al., 2017)or a lower dose (0.5mg/kg) of METH (Dobbs and Cunningham, 2014). We attribute this discrepancy to an effect of age, with adolescent C57Bl/6 males being less sensitive to 1mg/kg of METH than adults. This suggestion may be counterintuitive, as other CPP studies in rats (Zakharova et al., 2009) and a different mouse strain (Arc Swiss) demonstrate that adolescents are more sensitive to the rewarding effects of METH than adults (Cullity et al., 2021), depending on the dose used (Cullity, 2021). However, in the C57Bl/6 mouse strain, sensitivity to METH is not always stronger in adolescents. For example, METH has a greater effect on locomotion in adult than adolescent C57Bl/6 males, with opposite effects found at higher doses (Ortman et al., 2021). Adolescent C57Bl6 mice are less susceptible than adults to the neurotoxic effects of METH in the striatum (Miller et al., 2000). Furthermore, 2mg/kg of METH, a dose that is double that used in our study, leads to conditioned place aversion in a subset of adult C57Bl/6 males (Lominac et al., 2016; Szumlinski et al., 2017), an effect that has not been described in adolescent C57Bl/6 males (Buck et al., 2017). Adults may therefore be more sensitive to the aversive properties of METH than adolescents. Collectively, these results indicate that the effects of age on METH response are complex and dependent on both the strain and the dose that is used.

Sex differences in response to METH have been demonstrated in numerous rodent studies. For example, in rat self-administration studies, females escalate use faster and administer more METH than males (Kucerova et al., 2009; Roth and Carroll, 2004; Reichel et al., 2012). Female rats also exhibit more METH-seeking behavior during reinstatement than males (Cox et al., 2013; Reichel et al., 2012). In C57BL/6 mice, there are no sex differences in voluntary METH intake, but anticipatory mobility is higher in females than males (Avila et al., 2021). Furthermore, METH induces stronger CPP in adult C57Bl/6 females than males (Chen et al., 2003), similar to the sex difference we report in adolescent mice of this strain. Studies in other mouse strains demonstrate that METH-treated Arc Swiss females exhibit less conditioned place aversion than males (Cullity et al., 2021) and METH-treat BALB/c females exhibit a bigger increase in locomotor activity than males (Ohia-Nwoko et al., 2017). Together, these findings indicate that female-specific biological mechanisms can enhance the rewarding and stimulating effects of METH, with our results in C57Bl/6 mice indicating that these mechanisms may be developed as early as adolescence. However, our findings in 129Sv/Ev mice demonstrate that these sex differences are not maintained across all mouse strains.

The two mouse strains used in our study are widely used in the generation of knockout mice (Barnabei et al., 2010; Seong et al., 2004) and are known to differ in their physiology and behavior. Compared to the C57Bl/6 strain, 129Sv/Ev mice are less active (van Bogaert et al., 2006; Kelly et al., 1998; Good and Radcliffe, 2011), more susceptible to the effects of chronic stress (Aubry et al., 2019), and more anxious (Rodgers et al., 2002; Homanics et al., 1999). Our finding that adolescent 129Sv/Ev mice avoided the light compartment more than C57Bl/6 mice during preconditioning is consistent with higher baseline levels of anxiety in this strain. The 129Sv/Ev strain also has a higher baseline body temperature and is more sensitive than C57Bl/6 mice to the anxiolytic diazepam during the stress-induced hyperthermia test (van Bogaert et al., 2006). Unlike the C57Bl/6 strain, far fewer studies have tested the rewarding effects of drugs of abuse in 129 mice, most of which used 129 substrains that differ behaviorally from the 129Sv/Ev mice used here (Cook et al., 2002). These studies show that experimental conditions leading to cocaine self-administration or cocaine-induced CPP in C57Bl/6 mice, fail to do so in 129/OlaHsd or 129/SvJ mice, respectively (Miner, 1997; Kuzmin and Johansson, 2000). Similarly, the standard protocol used to detect morphine-induced CPP in C57Bl/6 mice is not sufficient to induce CPP in 129/SvJ mice (Dockstader and van der Kooy, 2001). These studies, all of which used adult males, indicate that 129s tend to be less sensitive to the rewarding effects of drugs of abuse than C57s, results that are seemingly the opposite of what we found in adolescent males. In our study, METH induced CPP and increased neural activity in 129Sv/Ev males, but not C57Bl/6 males. It is difficult to reconcile this discrepancy using published literature, as there are no previous studies that we are aware of that tested the rewarding effects of METH in any 129 substrain. Given that our study used adolescents instead of adults, one possibility is that there are strain differences in the development of the adolescent dopamine system that increase sensitivity to the rewarding effects of METH in 129Sv/Ev males and reduces sensitivity in C57Bl/6 males. It may be that response to METH and other drugs of abuse changes with further age-related development of the brain, resulting in a decrease in 129 sensitivity in adults, as previously reported. Future studies characterizing the development of the dopamine system in reward circuits of 129Sv/Ev and C57Bl/6 mice may provide insight into some of the mechanisms mediating the strain differences we report. Furthermore, understanding sex differences in these circuits may reveal why we found the opposite effect of METH on CPP in adolescent females (i.e. higher sensitivity in C57Bl/6 than 129Sv/Ev females).

Strain differences in the effects of psychostimulants on locomotor activity have been widely reported, with comparisons between C57Bl/6 mice and 129 substrains leading to mixed results. For example, cocaine has been shown to either increase locomotion similarly in both strains (Miner, 1997) or to stimulate locomotion in C57s only (Schlussman et al., 1998). For amphetamine, the stimulating effects on locomotion and dopamine efflux in the striatum are stronger in C57Bl/6 than 129S2/SvHsd mice (Chen et al., 2007). Similarly, we found that the first injection of METH led to a robust increase in locomotor activity in C57Bl/6 mice of both sexes, and a modest, but significant, increase in 129Sv/Ev females, but not males. These effects may be attributed to strain differences in the metabolism of METH in the striatum, which has been shown to be faster in 129Sv/Ev than C57Bl/6 males when administered at the same dose used in our study (Good and Radcliffe, 2011). Interestingly, we found that the stimulating effect of METH on locomotion was enhanced after the fourth injection in 129Sv/Ev males only, indicating that 1mg/kg is sufficient to induce both behavioral sensitization and CPP in this group. Even though C57Bl/6 females also exhibited CPP, they did not show sensitization, which may have required administration of a higher METH dose. Indeed, different threshold doses of amphetamine have been shown to induce CPP and sensitization in rats, with the latter requiring a higher dose (Rademacher et al., 2006).

We explored the neural substrates of METH-induced CPP in adolescent mice using c-Fos as a marker of neural activity. We found that mice exhibiting CPP had upregulated c-Fos protein expression in the NAc and hippocampus, results that are consistent with the known roles of both brain regions in conditioned reward. For example, CPP can be induced by local infusions of amphetamine into the NAc (Carr and White, 1983) and impaired by lesions of the hippocampus (Ito et al., 2006). Similar to our findings, the number of c-Fos+ cells in the NAc and CA1 of adult male rats is increased by amphetamine-CPP (Rademacher et al., 2006), indicating that the neural basis of CPP may be maintained across age, sex and species. Importantly, we found that neural activity in the NAc and CA1 is correlated in groups that exhibited METH-CPP, suggesting that communication between these brain regions is increased during retrieval of the conditioned memory. In line with these findings, a recent study showed that cocaine conditioning strengthens ventral CA1-to-NAc synapses, leading to the formation of an ‘engram circuit’ that stores CPP memory (Zhou et al., 2019). Similar processes may underlie METH-CPP in adolescent mice. Surprisingly, we found that females, but not males, of both genotypes had substantially more c-Fos+ cells in the NAc shell than the NAc core, regardless of drug exposure. The NAc shell has been implicated in the consolidation of memory for emotionally arousing events (Kerfoot and Williams, 2018), raising the possibility that females experience the non-drug related learning the occurs during the CPP procedure as more aversive than males. Alternatively, females may have more neural activity in the NAc shell than the NAc core at baseline. Although sex differences in NAc neuroanatomy (Forlano and Woolley, 2010; Wissman et al., 2011) and the regulation of dopamine in the NAc (Yoest et al., 2019) have been reported, we are not aware of other studies demonstrating sex differences in relative levels of neural activity across NAc subregions, particularly in the absence of drug.

To our knowledge, this study is the first to not only compare the effects of METH on CPP in 129Sv/Ev and C57Bl/6 mice, but to test the rewarding effects of METH in any 129 substrain. By administering METH during adolescence, our findings reveal strain differences in METH sensitivity during a critical period of development when the brain is undergoing ongoing maturation. In line with known sex differences in addiction-related behaviors, we report sex differences in the behavioral and neural effects of METH, although they were the opposite in each strain. This novel finding suggests that METH response may be determined by an interaction between sex-specific and genetic mechanisms. Finally, our finding that the expression of METH-CPP is associated with increased communication between the NAc and CA1 indicates that a NAc-CA1 circuit may mediate the storage and retrieval of METH CPP memory in adolescent mice. This study will inform future addiction studies in mouse models targeting mechanisms that differ across sex and strain.

## Acknowledgements

We thank Sadiyah Hanif for assistance with tissue processing. We would also like to acknowledge Barbara Wolin, Sonia Acevedo and other members of the animal care staff at Hunter College for excellent care of the animals.

## Author contributions

LNS, GM, and NSB wrote the manuscript. ABT and RML administered the drug and conducted the CPP experiments. NSB conducted all tissue processing. LNS conducted all cell quantification. LNS, ABT, RML, GM, and IB analyzed the behavioral data. LNS and NSB conducted final statistical analyses. All authors approved the manuscript for submission.

## Data availability

The raw data supporting the conclusions of this manuscript will be made available upon reasonable request to the corresponding author.

## Supplemental material

Supplemental material for this article is available online.

