## Supplementary Figure 1 for "Methamphetamine Conditioned Place Preference in Adolescent Mice: Interaction Between Sex and Strain"

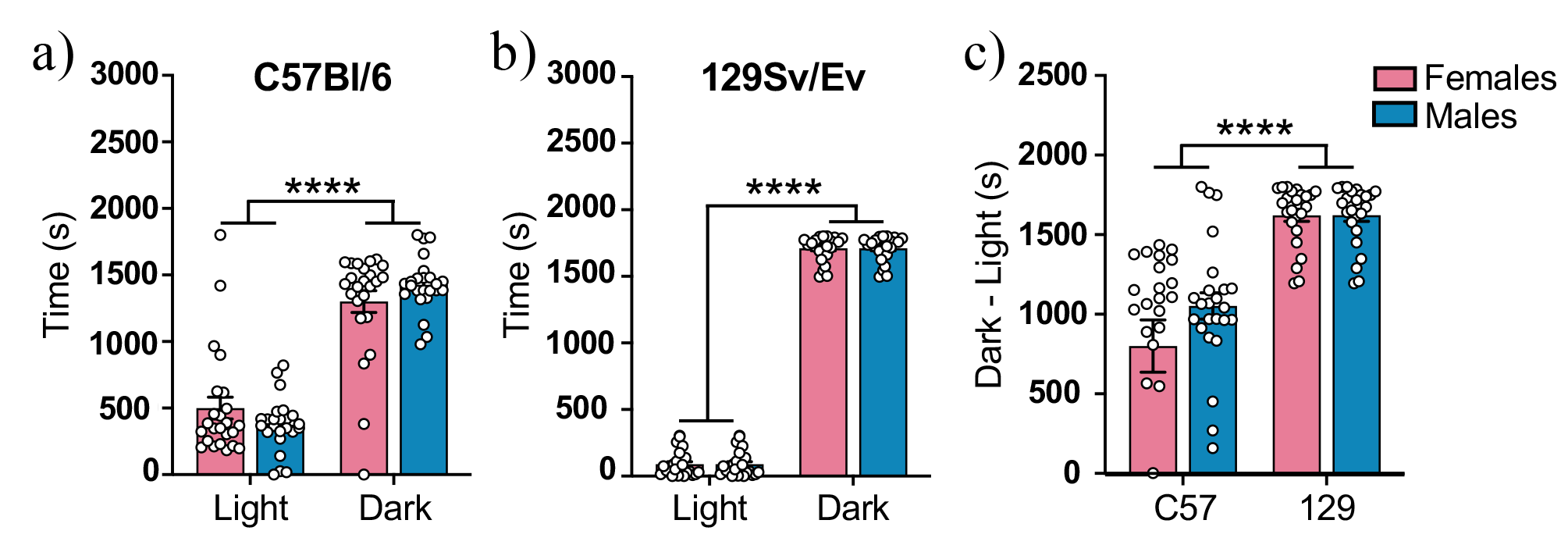

**Prior to conditioning, mice of both strains and sexes exhibited an innate preference for the dark side.** Time that **(a)** C57Bl/6 and **(b)** 129Sv/Ev male and female mice spent in each compartment during the preconditioning session. **(c)** Difference in time mice spent in the light and dark compartments, with higher numbers indicating stronger preference for the dark compartment. C57Bl/6 females (n=24), C57Bl/5 males (n=24), 129Sv/Ev females (n=24), 129Sv/Ev males (n=24). Data represent mean ± SEM. ****p<0.0001.
